# Evaluation of Immunopharmacological Efficacy of Live *Leishmania donovani* Overexpressing Ld_ζ1_domain_ for Protection Against Experimental Human Visceral Leishmaniasis

**DOI:** 10.1101/2024.12.30.630021

**Authors:** Ruby Bansal, Sadat Shafi, Prachi Garg, Aakriti Srivastava, Swati Garg, Neha Jha, Jhalak Singhal, Gajala Deethamvali Ghouse Peer, Ramendra Pati Pandey, Subhajit Basu, Shailja Singh

## Abstract

**Objective:** To evaluate the efficacy and immunogenicity of the zeta domain over-expressing *Leishmania donovani* (Ld_ζ1_domain_) as a vaccination candidate against visceral leishmaniasis (VL).

**Methods:** In this study, *Leishmania* overexpressor Ld_ζ1_domain_ (OE) were transformed by electroporation using a GFP-tagged Ld_ζ1_domain_ recombinant plasmid. The resulting overexpressing cells were analysed *in vitro* to assess their growth dynamics and infectivity. We also investigated the immune-protective potential of these overexpressor in a mouse model challenged with *Leishmania donovani*. The immune response, including Th1 and Th2 pathways, was thoroughly characterized using RT-PCR and ELISA assays. In addition, the study conducted a thorough evaluation of the mouse’s spleen and liver parasites, as well as quantitative evaluation of tissue pathological changes.

**Results:** Ld_ζ1 _domain_ (OE) parasites exhibited significantly lower viability and replication rates than WT parasites, and *in vivo* studies showed that mice immunized with the Ld_ζ1(OE) _domain_ had lower parasite numbers than mice infected with LdWT. Spleen and liver showed significant histological changes suggestive of protection. Parasite burden in the spleen and liver of vaccinated mice were significantly reduced. The immune response showed increased IFN-γ levels and lower IL-10 production, resulting in a greater IFN-γ/IL-10 ratio, indicating parasite elimination. The vaccination also caused a significant IgG humoral response and increased nitric oxide production in immunized mice.

**Conclusion:** Our findings demonstrated that overexpressing the zeta toxin resulted in controlled parasite attenuation, lowering pathogenicity while retaining immunogenic features. Our work established the zeta over-expressor’s protective efficacy, immunogenicity, and proliferation in response to a *Leishmania* challenge *in vitro* and *in vivo*. This preliminary prototype study suggested that Ld_ζ1_domain_ (OE) parasites may be suitable for developing an attenuated vaccine against leishmaniasis.

**Graphical representation:** 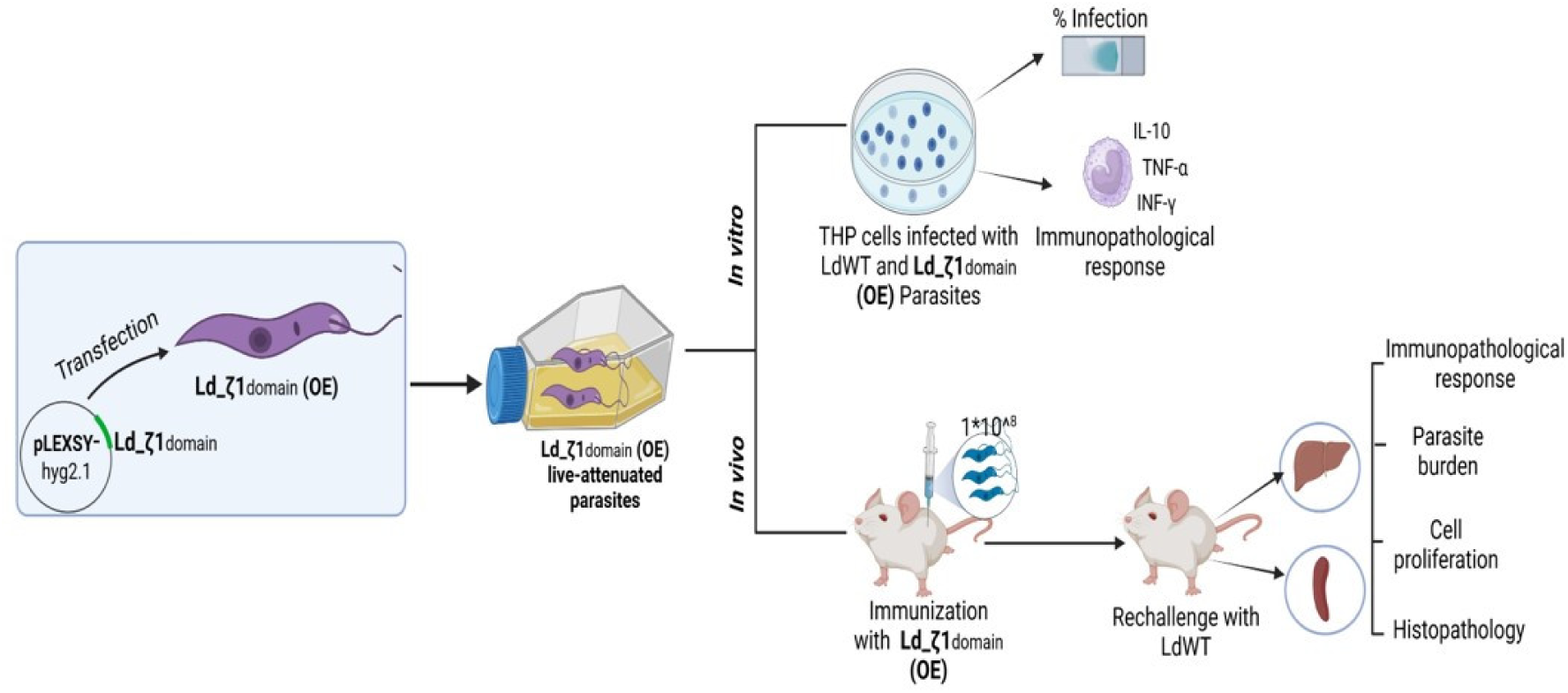

A schematic representation on the protective effectiveness, immunogenicity, and proliferation of the zeta over-expressor in response to *Leishmania* challenge *in vitro* and *in vivo* model.

## 1. Introduction

Leishmaniasis, a neglected protozoal parasitic disease, is caused by an infection with the obligatory intracellular parasite *Leishmania* and transmitted by infected sand flies. Leishmaniasis has several clinical presentations, including Cutaneous Leishmaniasis (CL), Mucocutaneous Leishmaniasis (MCL), and Visceral Leishmaniasis (VL) [1]. VL often known as kala azar in India, is the most severe type induced by *Leishmania donovani* (Ld) infection. VL spreads to internal organs such as the spleen and bone marrow, leading in around 30,000 new cases every year (World Health Organization, 2023), with substantial death rates in the absence of adequate treatment [2].Current leishmaniasis treatment relies on pentavalent antimonial, miltefosine and amphotericin B (AmB) [3]. However, the increasing resistance to first-line treatments and the adverse effects of second-line treatments underscore the pressing need for an effective therapeutic strategy and vaccine against this disease.

Although no licensed vaccine exists for any form of leishmaniasis, however numerous antileishmanial vaccine formulations including live attenuated *Leishmania* vaccines, genetically modified strains, heat-killed parasite, DNA and subunits have been tested in animal models, but few advance beyond experimental stages [4].

Live attenuated *Leishmania* parasites, non-pathogenic and mimicking the wild-type strains’ antigenicity, emerge as promising candidates for evoking immunologic memory [5]. Parasites have been suppressed using a variety of methods, including gamma irradiation and genetic alterations [6]. Consequently, several experimental live-attenuated vaccines have been generated by deleting or knocking out a range of genes. Recent investigations show that centrin-deficient *L. mexicana* and *L. donovani* strains protect against CL and VL, respectively [7,8]. Centrin is a cytoskeletal protein required for eukaryotic cell division but exclusively for intracellular amastigote multiplication in *Leishmania*.

Previously, many studies have discovered that the inoculation with an *L. donovani* strain lacking the biopterin transporter (BT1) protected mice [9] against *Leishmania* challenge. Additionally, *L. donovani* mutants lacking the amastigote-specific p27 gene have been demonstrated to be safe and provide cross-protection against *L. major* [10]. Partial knockout parasites for the A2-A2rel gene cluster in *L. donovani* [11] and the SIR2 gene in *L. infantum* [12] were also shown to protect BALB/c mice against virulent challenge. However, the security of such variants cannot be assured because they still include a wild type allele that may cause disease. Therefore, it is crucial to establish attenuated lines which have a lower probability of reactivation.

In the current investigation, we generated genetically modified live-attenuated *Leishmania* parasites by overexpressing a zeta toxin, which is a vital component of the toxin-antitoxin (TA) system. Overexpression of a toxin lead to compromised and weakened parasites. These compromised organisms are capable to elicit an immune response without the risk of pathogenicity, offering a promising strategy for vaccine development. TA are naturally occurring substances with both bacterial and eukaryotic origins. Toxins prevent cell growth by interfering with vital biological functions, whereas antitoxins combat toxins. Stress reduces antitoxin expression, allowing toxins to disrupt cellular functions such as replication and transcription [13,14]. Their function is often associated to offensive or defensive reactions, which lead to the damage or destruction of the targeted organism [15]. Artificial activation of the plasmid-encoded TA system acted as an anti-bacterial strategy followed by the typical toxin-mediated cell death mechanism [16].

We previously demonstrated the existence of a prokaryotic like zeta toxin protein in *L*. *donovani* (Ld_ζ1) and its characterisation in promastigotes and a heterologous prokaryotic system, *Escherichia coli* [17]. According to published genome sequences, the zeta toxin of the type-II TA system of prokaryotes is one such protein that is thought to be present in the majority of kinetoplastid species. Ld_ζ1, a *Leishmania* zeta-toxin with similar UNAG and ATP-binding sites, exhibited ATP-binding and UNAG kinase activity. Ld_ζ1 is an active protein in *L. donovani*, which functions in growth regulation.

In the current study, we first cloned the Ld_ζ1_domain_ into a GFP-tagged *Leishmania*-specific vector and overexpressed it in *L. donovani* promastigotes by transfection to generate zeta domain over-expressor Ld_ζ1_domain_ (OE). We further characterized the Ld_ζ1_domain_ (OE) promastigotes and investigated the growth and infectivity of Ld_ζ1_domain_ (OE) in THP-1 cells in *vitro*. Additionally, our investigation examined the protective effectiveness, immunogenicity, and proliferation of the Ld_ζ1_domain_ (OE) in response to *Leishmania* challenge in a murine model. Moreover, we endeavoured to understand the immunological processes underlying the observed protective benefits.

Our findings attempted to study the potential of Ld_ζ1_domain_ (OE) in protection against VL and acted as a preliminary research effort to find a novel prophylactic strategy for combating leishmaniasis, addressing a huge worldwide health burden.

## 2. Materials & Methods

### 2.1 Animals and parasites

Male Swiss mice, aged four to six weeks, were procured from Jawaharlal Nehru University’s (JNU) Animal Facility. All animals were housed in a centralized facility under aseptic conditions. The wild-type *L. donovani* (LdWT) (MHOM/SD/62/1S) and *L. donovani* (Ld_ζ1_domain_) over-expressor promastigotes were cultured in liquid M199 with Hanks’ salts (Gibco, ThermoFischer Scientific, USA) culture medium augmented with 10% fetal bovine serum (FBS, Gibco, ThermoFischer Scientific, USA) and 1% Penicillin-Streptomycin (Gibco, ThermoFischer Scientific, USA) at 26°C. The animal experiments were carried out at Jawaharlal Nehru University’s Central Laboratory Animal Resources (CLAR) in New Delhi, India. The care and treatment of experimental animals was in compliance with the Institutional Animal Ethics Committee (IAEC) of JNU.

### 2.2 Growth curve of promastigotes & axenic amastigotes

The growth of *L. donovani* promastigotes and axenic amastigotes was analysed continuously over a 10-day period at 26°C and 37°C, respectively, in their specific media. Cells were seeded in 96-well plates (Nunc, Roskilde, Denmark) at a density of 4 x 10^4^ cells/mL, with the analysis focusing on exponentially growing populations. We successfully established axenic cultures of amastigote forms for both wild-type (WT) and Ld_ζ1_domain_ (OE) strains of *L. donovani.* These axenic amastigotes were maintained at 37°C with 5% CO_2_ in M199 medium at pH 5.5, containing Hanks’ salts and supplemented with 20% FBS, in 25-cm^2^ flasks [18].

### 2.3 Immunofluorescence assay

Promastigotes expressing the Ld_ζ1_domain_ were immobilized on poly-L-lysine-coated coverslips (Thermo Fisher Scientific, USA) to investigate zeta protein intracellular localization. The cells were fixed and permeabilized, then incubation with anti-Ld_ζ1_domain_ mouse sera (1:500) for 1 h at room temperature (RT). Following washing, the cells were incubated for 45 min with Alexa Fluor 488-conjugated goat anti-mouse IgG (H+L) antibody (Invitrogen, Thermo Fisher Scientific, USA). Nuclear and kinetoplast DNA were stained with DAPI (Invitrogen, Thermo Fisher Scientific, USA). Additionally, co-localization of the Ld_ζ1_domain_ was also examined using LysoTracker (Invitrogen, Thermo Fisher Scientific, USA), a dye specific for acidic organelles. The immunofluorescence staining of the parasites was observed using a fluorescence microscope (Nikon, Towa Optics (I) Pvt Ltd., New Delhi, India.

### 2.4 Real-time quantitative PCR

Total RNA was extracted from cells at the relevant time point using TRIzol reagent (Invitrogen, Grand Island, NY, USA), washed with 75% ethanol, and then reconstituted in RNase-free water. The PrimeScript 1st Strand cDNA Synthesis Kit (Takara Bio, USA) was used to synthesize cDNA from 1 µg of total RNA, following manufacturer instructions. Real-time PCRs were carried out using the Applied Biosystems Real-Time PCR (RT-PCR) System (ABI, Foster City, CA, USA) and PowerUp SYBR Green PCR Master Mix (Thermo Fisher Scientific, USA). Table 1 details the primer sequences utilized in the RT-PCR assays.

**Table 1:**
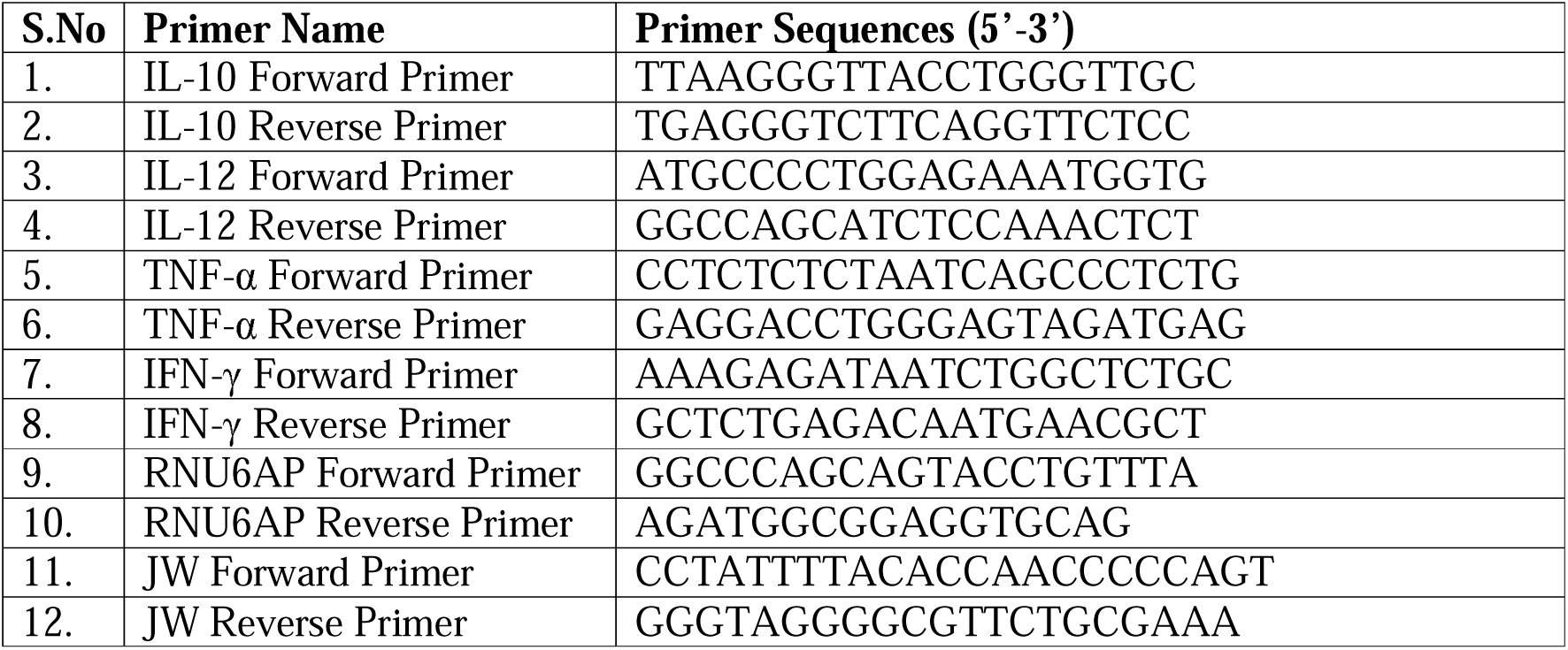
Primer sequences used for Real time qPCR analysis:

To determine cytokine levels, RNA was isolated from respective cells and levels of IFN-γ, TNF-α, and IL-10 were quantified by RT-PCR after 48 h of incubation using specific primers. Amplification of RNU6AP (RNA, U6 small nuclear 1; Human THP-1 macrophages) and GAPDH (Mouse) served as internal controls for normalization. The following temperature cycling conditions were used: 50°C for 2 min, accompanied by 40 cycles of 95°C for 15 s, 60°C for 30 s, and 72°C for 1 min. Gene expression levels were measured using the 2^−ΔΔCt^ method and calibrated against the housekeeping gene. Melting curves were created along with mean C_t_ values to corroborate the specificity of the PCR products. Data analysis was analyzed using GraphPad software.

### 2.5 Human macrophage infection

THP-1 cells were obtained from the National Centre for Cell Science (NCCS, Pune, India). The cells were cultured in RPMI-1640 medium (Gibco-Life Technologies, USA) enriched with 10% heat-inactivated fetal bovine serum, 10 mM pyruvate, 10 mM L-glutamine, and 1% penicillin-streptomycin. THP-1 monocytes were allowed to differentiate into macrophages by using 100 ng/mL phorbol 12-myristate 13-acetate (PMA, Sigma-Aldrich, USA) [19]. After differentiation, macrophages were then infected with stationary phase LdWT and Ld_ζ1_domain_ (OE) promastigotes at a 20:1 parasite-to-macrophage ratio, while uninfected macrophages served as controls. After 6 h incubation at 37°C in 5% CO_2_, free extracellular parasites were washed away with incomplete RPMI medium, and the cultures were further incubated in media for an additional 24 h and 48 h. The culture medium was then removed, and the infected macrophages were fixed with 100% methanol for 2 min. This was followed by Giemsa staining for 3 min, subsequent washing with distilled water, and air drying. The percentage of infected macrophages was calculated by counting at least 100 macrophages per sample under a microscope. Results are shown as mean ± SEM from three separate counts for each infection.

### 2.6 Immunization schedules of mice, parasite burden estimation and parasitized splenic cell isolation

After acclimatization, mice were segregated into three groups: (a) control uninfected group, (b) unimmunized group but challenged with LdWT parasites, and (c) immunized group with Ld_ζ1_domain_ (OE) parasites and challenged with LdWT. For immunization, 1×10^8^ total stationary-phase Ld_ζ1_domain_ (OE) parasites were used via intravenous injection into the tail. After six weeks, a subset of mice from each group was euthanized, while the remaining mice were treated with 2.5 mg/kg amphotericin B (AmB) (Sigma Aldrich, USA) for 5 days to clear the infection. One week after post-clearance, or eight weeks post-immunization, the mice were rechallenged via tail vein injection with 1×10^8^ stationary-phase LdWT promastigotes, followed by termination of the experiment after two weeks. At each experimental time point (6^th^, 7^th^, and 10^th^ week post-immunization), four inoculated mice from each group were euthanized. Parasite burdens in the spleen and liver were determined by RT-PCR and haematoxylin and eosin (H & E) staining. In each study, four mice were used per group per time point.

### 2.7 Histopathological studies

Spleen and liver tissues were collected post-euthanasia at each experimental time point and fixed in 4% paraformaldehyde to preserve their structure. These tissues were then processed, sectioned, and stained with hematoxylin and eosin to highlight cellular details. Under a Zeiss light microscope, the sections were examined for the presence of parasites and inflammatory infiltrates, providing valuable insights into the progression of infections and the associated immune responses.

### 2.8 Serological studies and analysis

IgG-specific antibody responses were quantified using standard ELISA. Blood was collected from mice through the tail vein, and serum was separated by centrifugation. A 96-well plate was coated with 80 μg/mL of freeze-thawed *Leishmania donovani* antigen (FTA) and incubated overnight at 4°C. Following washing, plate was blocked and treated with serum samples in duplicates and underwent incubation with HRP-conjugated anti-mouse IgG antibodies (1:1000 dilution, Sigma-Aldrich, USA). The tetramethylbenzidine (TMB) (Sigma-Aldrich, USA) substrate was added, and the reaction was terminated using 5% sulfuric acid. Absorbance was measured at a wavelength of 450 nm via a Molecular Devices SpectraMax This approach offers a specific and sensitive evaluation of the humoral immune response to the *Leishmania donovani* antigen.

### 2.9 Cell proliferation and cytokine assays

Spleens were aseptically harvested from experimental mice at designated time points, and single-cell suspensions were meticulously prepared in RPMI-1640 medium supplemented with penicillin G (100 U/ml), streptomycin sulfate (100 mg/ml), and 10% heat-inactivated FBS. Following two thorough washes, splenocytes were resuspended in culture medium and seeded in triplicate into 96-well flat-bottom plates at a density of 2 x10^5^ cells per well in a total volume of 200 μl of complete medium. The cells were stimulated with freeze-thaw *Leishmania* antigen (FAg; 80 μg/ml) and incubated at 37°C in a humidified chamber with 5% CO_2_. After 48 h, supernatants were carefully collected for cytokine analysis. Levels of mouse TNF-α (430904), IFN-γ (430804), and IL-10 (431414) were quantified using BioLegend ELISA kits (San Diego, CA, USA), strictly adhering to the manufacturer’s protocols.

### 2.10 Nitric Oxide (NO) quantification

Supernatants collected from freeze-thaw *Leishmania* antigen stimulated splenocytes were used to determine nitric oxide (NO) production. NO levels, as measured by nitrite/nitrate concentrations, were determined using the Griess Reaction Kit (Sigma Aldrich, USA). This assay includes converting nitrate to nitrite, followed by a colorimetric reaction using sulfanilamide and N-(1-naphthyl) ethylenediamine dihydrochloride, which produces a detectable azo dye. The method was carried out in accordance with the manufacturer’s instructions to ensure precise and reliable outcomes.

### 2.11 Ethics statement

The animal protocol for this research has been authorized by Jawaharlal Nehru University’s Institutional Animal Ethics Committee (IAEC) in New Delhi, India. Animal studies were done at the Central Laboratory of Animal Resources (CLAR), JNU.

### 2.12 Statistical analysis

The student’s t test was used with Graph Pad Prism 7.0 software to detect statistical differences between group means. The figure legends describe the statistical tests and their associated significance values. P-values <0.05 indicated statistical significance. p-Values were denoted as *p < 0.05, **p < 0.01, ***p < 0.001, and ****p < 0.0001.

## 3. Results

### 3.1 Generation and characterization of Ld_ζ 1_domain_ over-expressor

Zeta toxins, part of Type-II toxin-antitoxin systems, are underexplored in eukaryotes. We identified three Zeta toxin-like proteins in *Leishmania donovani.* One variant, LdBPK_341740.1, is a 1014 amino acid protein (∼111.5 kDa) featuring a 118 amino acid zeta domain (Ld_ζ1_domain_) **[Figure 1A]** and a P-loop nucleoside triphosphate hydrolase domain. This unique structure suggests a different functional role in *L. donovani* compared to prokaryotic counterparts.

**Figure 1:**
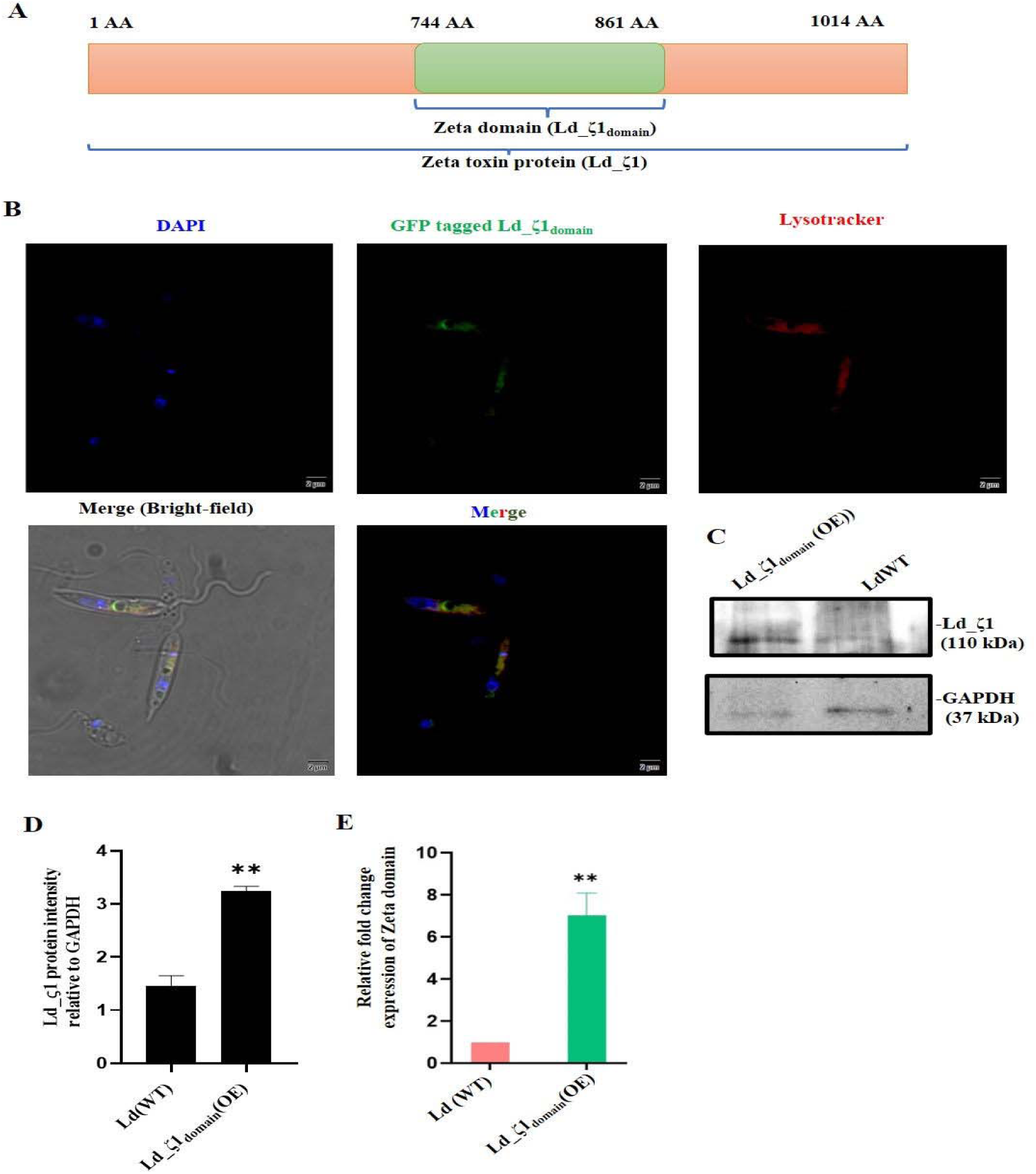
**Characterization of GFP tagged Ld_**ζ**1_domain_ (OE) promastigotes A**) Analysis of the *Leishmania donovani* genome revealed a novel Zeta toxin-like protein, designated as Ld_ζ1. This protein contains a 118-amino acid zeta domain, referred to as the Ld_ζ1_domain_. **B**) Immunofluorescence microscopy was utilized to assess the expression of Ld_ζ1 in fixed promastigotes. Promastigotes were probed with sera raised in mice against the Ld_ζ1_domain_. The secondary antibody used was conjugated with Alexa 488, allowing visualization. Co- localization of the Ld_ζ1_domain_ was examined by staining with LysoTracker, a dye that identifies acidic organelles, to validate the presence of the protein in acidic organelles associated with autophagy. The scale bar in the images represents 10 µm **C**) Western blot analysis was used to evaluate Zeta protein expression in *Leishmania* cell lysate (40 µg) by using zeta domain antibodies raised in mice. GAPDH was used to ensure internal normalization. **D**) The relative intensity of the protein bands that belong to zeta protein was adjusted to GAPDH and assessed using ImageJ software.**E**) RT-PCR analysis to demonstrate the presence of the Ld_ζ1 gene in *L. donovani* promastigotes. The process involved isolation of RNA from promastigotes, synthesizing cDNA, and amplifying the Ld_ζ1 gene to confirm mRNA expression.

To elucidate the subcellular localization and function of the Ld_ζ1 protein in *Leishmania donovani*, we employed a protein overexpression strategy in promastigotes by overexpress a GFP-tagged Ld_ζ1_domain_ in *Leishmania*. The Ld_ζ1_domain_ was cloned into pLEXSY-hyg2.1 vector and transiently transfected into promastigotes to produce target protein overexpression. Following hygromycin B selection of transfectants, we validated the expression of these constructs in promastigotes using GFP fluorescence. The GFP-tagged Ld_ζ1_domain_ was found localized in vacuole-like structures. The co-localization of Ld_ζ1_domain_ was examined by staining with lysotracker, a dye that identifies acidic organelles. The co-localization of Ld_ζ1_domain_ was examined by staining with lysotracker, a dye that identifies acidic organelles. The co-staining of GFP fluorescence of Ld_ζ1_domain_ with lysotracker confirms that the domain is positioned in a vacuole-like structure linked to programmed cell death [**Figure 1B**].Zeta protein expression was measured by western blotting with antibodies generated against the zeta domain in mice, which showed higher levels of zeta protein [**Figure 1C]**. GAPDH was used as an internal normalisation control. The relative intensity of the protein bands corresponding to zeta protein was normalised and quantified against GAPDH with ImageJ software [**Figure 1D].** To demonstrate the presence of the putative Ld_ζ1 gene in *L. donovani* promastigotes, we amplified a segment of the Ld_ζ1_domain_ using *L. donovani* cDNA as a template, resulting in a sixfold increase in Ld_ζ1_domain_ expression at the transcript level [**Figure 1E]**.

### 3.2 Growth kinetics and infectivity of Ld_**ζ**1_domain_ (OE)

We investigated the growth dynamics of promastigotes and amastigotes overexpressing the Ld_ζ1_domain_ to determine if it has toxicity effects in *Leishmania donovani*, potentially triggering parasite attenuation. During the early log phase, there were no significant variations in growth patterns between the overexpressing promastigotes and amastigotes of Ld_ζ1_domain_ (OE) and the WT controls. However, in the exponential growth phase, starting after 5 days, amastigotes overexpressing the Ld_ζ1_domain_ witnessed a substantial delay in growth compared to WT. Overexpression of the Ld_ζ1_domain_ inhibits development of *L. donovani* amastigotes **[Figure 2A]**. We observed similar growth kinetics between LdWT and Ld_ζ1_domain_ (OE) promastigotes. However, Ld_ζ1_domain_ (OE) amastigotes exhibited a notable growth defect starting at day 6 compared to LdWT amastigotes.

**Figure 2:**
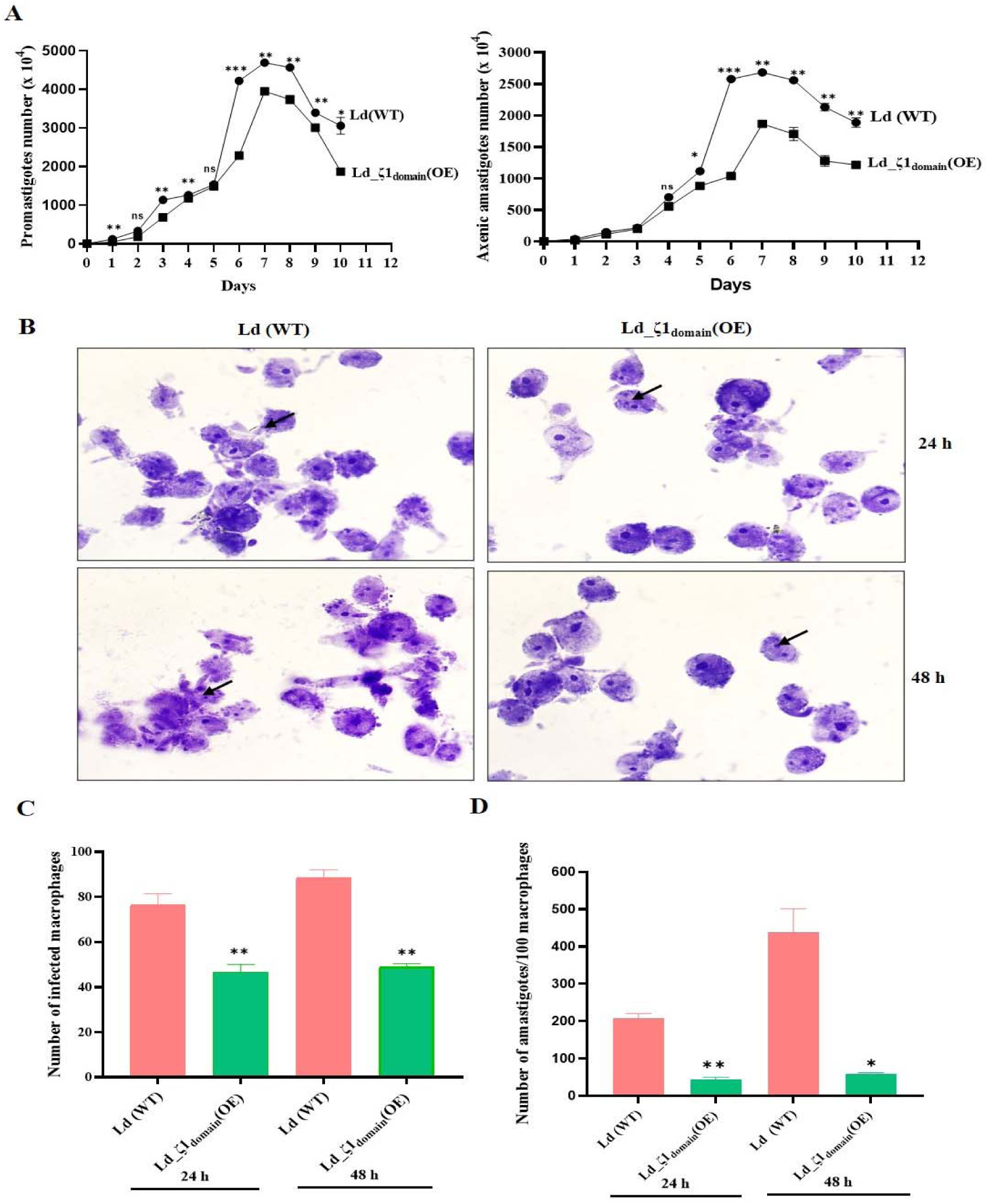
**Growth kinetics and infectivity of Ld_**ζ**1_domain_ (OE). A)** Growth curves of WT and Ld_ζ1_domain_ (OE) promastigotes and amastigotes. The growth pattern of *L. donovani* promastigotes and axenic amastigotes was analysed continuously over a 10-day period at 26°C and 37°C respectively, in their specific media. Cells were seeded in 96-well plates at a density of 4 x 10^4^ cells/mL. **B)** Giemsa-stained micrograph showing human macrophage cell line (THP-1) infected with WT and Ld_ζ1_domain_ (OE) promastigotes. Macrophages showing intracellular parasite converted to amastigotes after internalization (100x; indicated by arrowheads). **C**) Percentage of infected macrophages was measures after 24 h and 48 h **D**) The quantity of intracellular parasites per 100 macrophages was determined at 24 h and 48 h post infection. The bars show the mean ± SD of four independent tests conducted in triplicate. An unpaired two-tailed Student’s t-test was used to compare statistical significance at each time point. P-values < 0.05 were regarded as significant. In all panels, * denotes P ≤ 0.05, ** P ≤ 0.01, and *** P ≤ 0.001.

In this study, the rate of infection was estimated using giemsa-stained slides by light microscopy **[Figure 2B]**. The results revealed that macrophages infected with Ld_ζ1_domain_ (OE) parasites exhibited a reduction in the number of infected macrophages to 30-40% at 24 h and 48 h post infection **[Figure 2C]**. The quantity of intracellular parasites per 100 macrophages reduced by 75% and 85% after 24 and 48 h, respectively, as compared to macrophages infected with WT parasites **[Figure 2D]**. Ld_ζ1_domain_ (OE) parasites had considerably lower survival and replication rates than WT parasites.

### 3.3 Ld_**ζ**1_domain_ (OE) enhanced immune response and parasite clearance in THP-1 cells

Additionally, we evaluated the parasitic burden using quantitative RT-PCR to assess the fold change expression of the *Leishmania*-specific kinetoplast gene (JW) in THP-1 cells infected with WT and Ld_ζ1_domain_ (OE) parasites. This investigation aimed to compare the capacity of Ld_ζ1_domain_ (OE) parasites to modulate responses in THPs *in vitro*. Our findings revealed that LdWT parasites exhibited heightened infection rates and a greater parasite load, quantified via RT-PCR, in comparison to Ld_ζ1_domain_ (OE) parasites. Remarkably, we observed a significant 7-fold reduction in infection in Ld_ζ1_domain_ (OE) infected macrophages compared to those infected with LdWT parasites **[Figure 3A]**. This suggested a potential for Ld_ζ1_domain_ to attenuate parasitic burden effectively.

**Figure 3:**
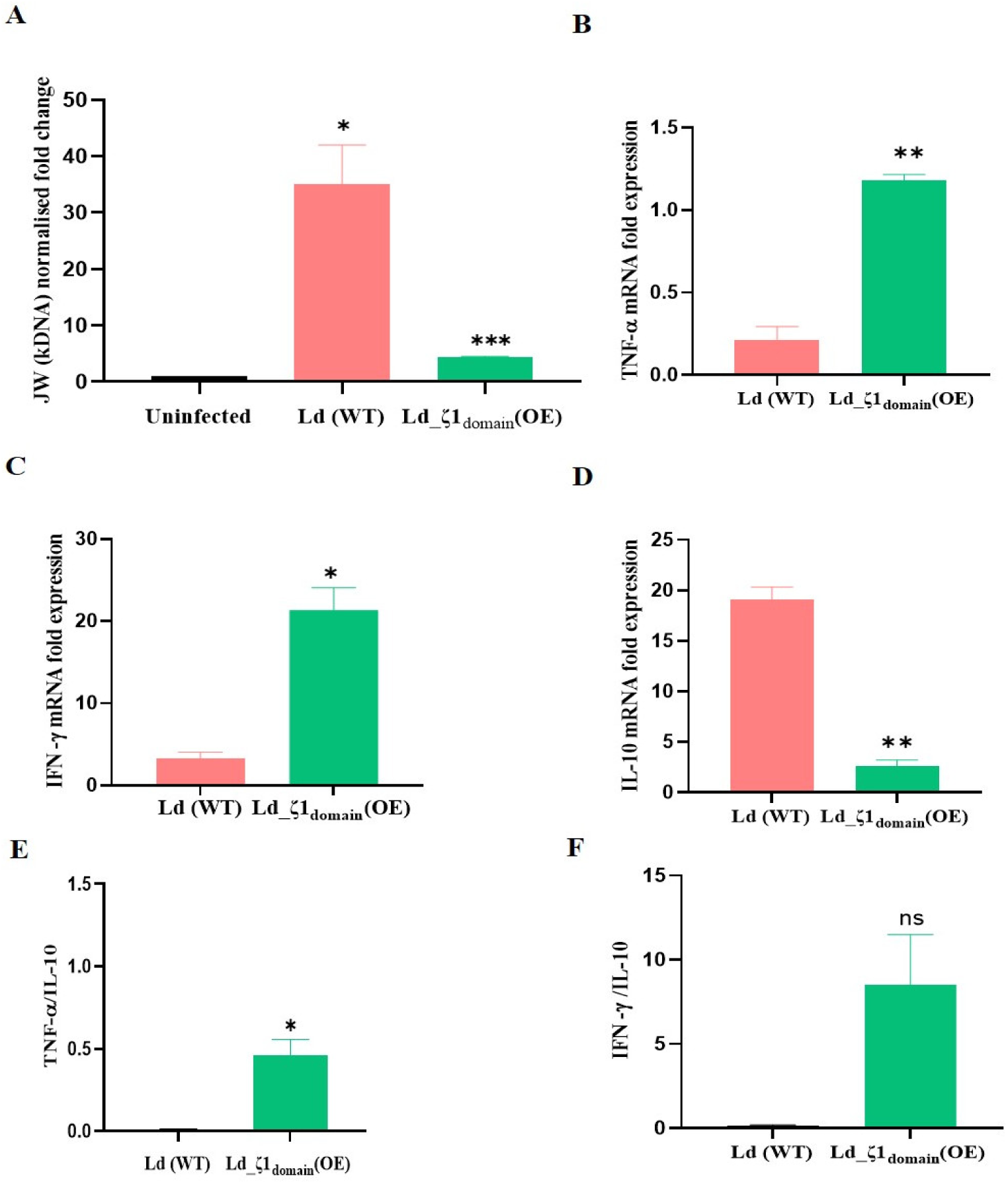
Protective innate immune responses *in vitro*: **A**) Ld_ζ1_domain_ (OE) parasites show diminished virulence in THP-1 macrophages. **B**) Pro-inflammatory TNF-α and **C**) IFN-γ & **D**) anti-inflammatory IL-10 cytokines production induced by WT and Ld_ζ1_domain_ (OE) in PMA-differentiated THP-1 cells. mRNA expression was determined by quantitative real-time PCR (24 h). **E**) & **F**) Ratio of TNFα/IL-10 and IFN-γ/IL-10 cytokines were also plotted which suggested the enhanced protection and clearance of parasites. Syber green RT-PCR was used to detect amastigote transcripts using kinetoplast specific primers, and the proportion of expression was standardized to RNU-6P. Results are shown as a percentage of untreated control (± SD) and are based on at least three separate studies. The unpaired two-tailed Student’s t-test was used to examine statistical significance at each time point. A P-value < 0.05 was considered significant. In all panels * represents P ≤ 0.05, ** represents P ≤ 0.01, and *** represents P ≤ 0.001

We also examined the mRNA expression of proinflammatory and anti-inflammatory cytokines in THP-1 cells infected with Ld_ζ1_domain_ (OE), contrasting them with LdWT parasitized macrophages. Notably, THP macrophages exhibited enhanced effector function in response to Ld_ζ1_domain_ (OE) infection *in vitro*.

Following a 24 h period, we observed a significant upregulation of TNF-α **[Figure 3B]** and IFN-γ **[Figure 3C]**, but not IL-10 **[Figure 3D]**, in both LdWT and Ld_ζ1_domain_ (OE) infections. Interestingly, Ld_ζ1_domain_ (OE) induced notably higher expression levels of TNF-α and IFN-γ, while concurrently displaying significantly lower levels of IL-10 in infected THP-1 cells. This reduced induction of Th2 cytokines led to elevated ratios of TNFα/IL-10 **[Figure 3E]** and IFN-γ/IL-10 **[Figure 3F]** cytokines, suggestive of enhanced protection and clearance of parasites.

### 3.4 Immunization with Ld_**ζ**1_domain_ (OE) confers protection to mice upon challenge with virulent *L. donovani*

The immunization strategy involving Ld_ζ1_domain_ (OE) in mice is outlined in **Figure 4A** in schematic form. This diagram delineates three distinct categories across specific time points: Immunization, Infection clearance, and Re-infection. Immunization represents the endpoints post the initial challenge, Infection clearance signifies data following AmB treatment, and Re-infection indicates the *Leishmania* challenge administered to the mice groups subsequent to AmB treatment. These categories meticulously delineate each experimental time point, namely the 6^th^, 7^th^, and 10^th^ week post-immunization.

**Figure 4:**
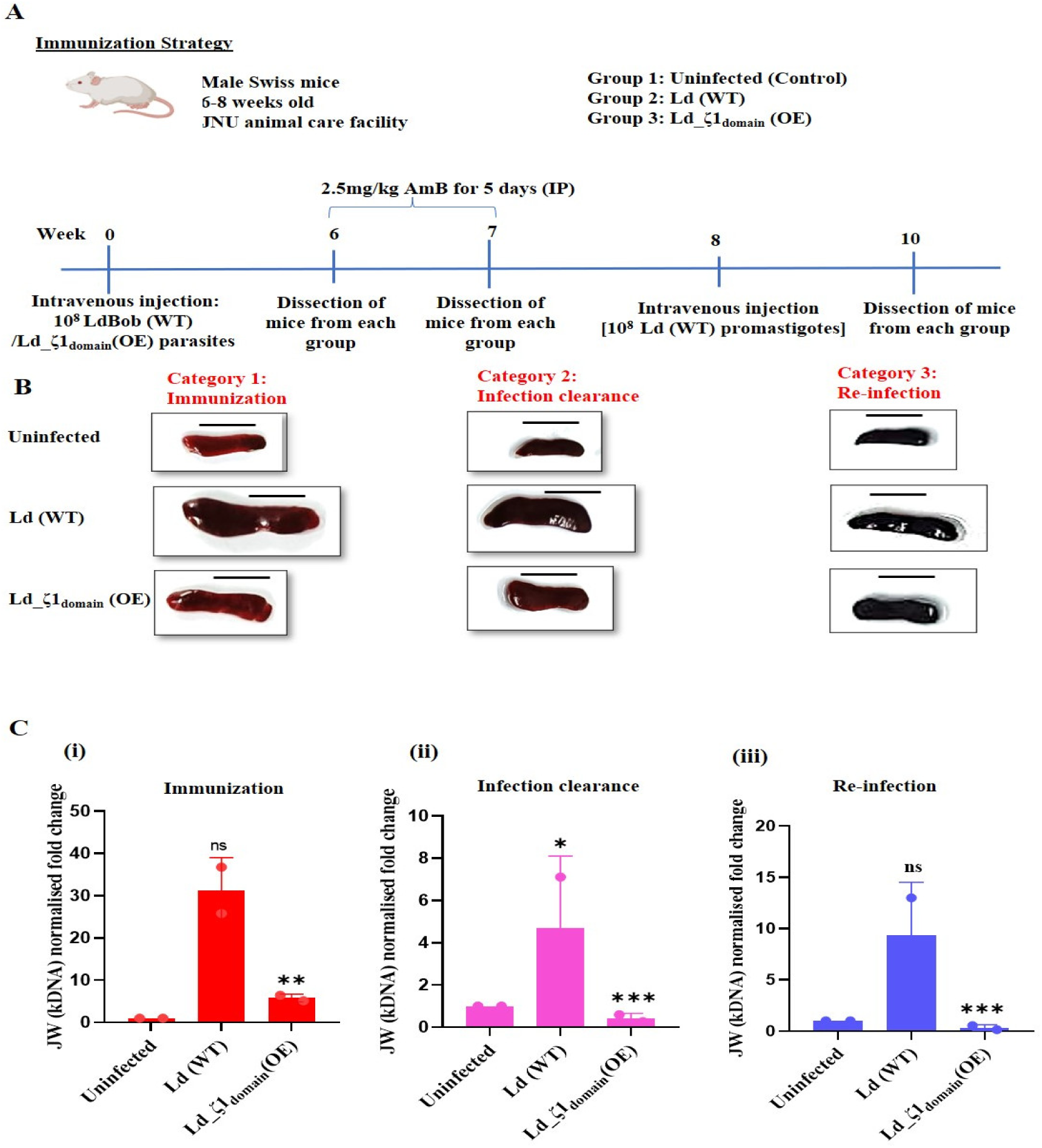
**Immunization with Ld_**ζ**1_domain_ (OE) confers protection to mice upon challenge with virulent *L. donovani.* A)** Schematic representation of immunization strategy used for *in vivo* studies. This diagram delineates three distinct categories across specific time points: **B**) Immunization, Infection clearance, and Re-infection. Four mice were used per group per time point. Comparison of the spleen size of Ld_ζ1_domain_ (OE) immunized animals to the uninfected group. Scale bar =1 cm **C)** RT-PCR analysis was performed at each point **i**) Immunization, **ii)** Infection clearance, and **iii)** Re-infection to determine the parasite burden in splenic tissue and to check infection level in immunized and unimmunized mice as done by checking JW expression which is housekeeping gene of the parasites.

Splenomegaly is a significant clinical symptom of VL. In experimental murine VL, there is a distinct evolution of splenomegaly concomitant reorganization of the spleen compartments [20]. The spleen size was consistently enlarged in WT-infected mice as compared to the mice group which were immunized with Ld_ζ1_domain_ (OE) parasites across all experimental conditions. Notably, the spleen size of Ld_ζ1_domain_ (OE) immunized animals remained similar to that of the uninfected group **[Figure 4B]**. To correspond with the observed splenic size disparities, we performed RT-PCR analysis to determine the parasite burden in splenic tissue. Mice infected with LdWT had a sixfold increased parasite burden in splenic tissues compared to Ld_ζ1_domain_ (OE) immunized mice soon after immunization **[Figure 4C (i)]**. At 7^th^ and 10^th^ week post-immunization also, LdWT infected mice had significantly greater parasitemia than Ld_ζ1_domain_ (OE) immunized mice **[Figure 4C(ii) & (iii)]**.

### 3.5 Comparative analysis of splenic and hepatic granuloma formation and parasite infection rates in Ld_**ζ**1_domain_ over-expressor versus WT infected mice

Histological disorganization of splenic architecture was observed across WT infected mice groups which has been associated with disease progression **[Figure 5A (i)]**. Granuloma examination in infected mice spleens (bar = 20 μm) revealed a higher infection rate in LdWT infected splenocytes while Ld_ζ1_domain_ (OE) infected splenocytes showed a 50% reduction in infection when parasitemia was quantified by determining the number of parasites per 100 splenocytes **[Figure 5A (ii)]**. Following amphotericin treatment, infection levels decreased in both groups **[Figure 5B (i) &(ii)]**. Upon reinfection, Ld_ζ1_domain_ (OE) immunized mice sustained a protective immune response, resulting in a substantially reduced parasite load in the spleen. This shows that vaccination caused long-lasting immunological memory, facilitating more effective management of the parasite than non-immunized controls **[Figure 5C (i) &(ii)]**.

**Figure 5:**
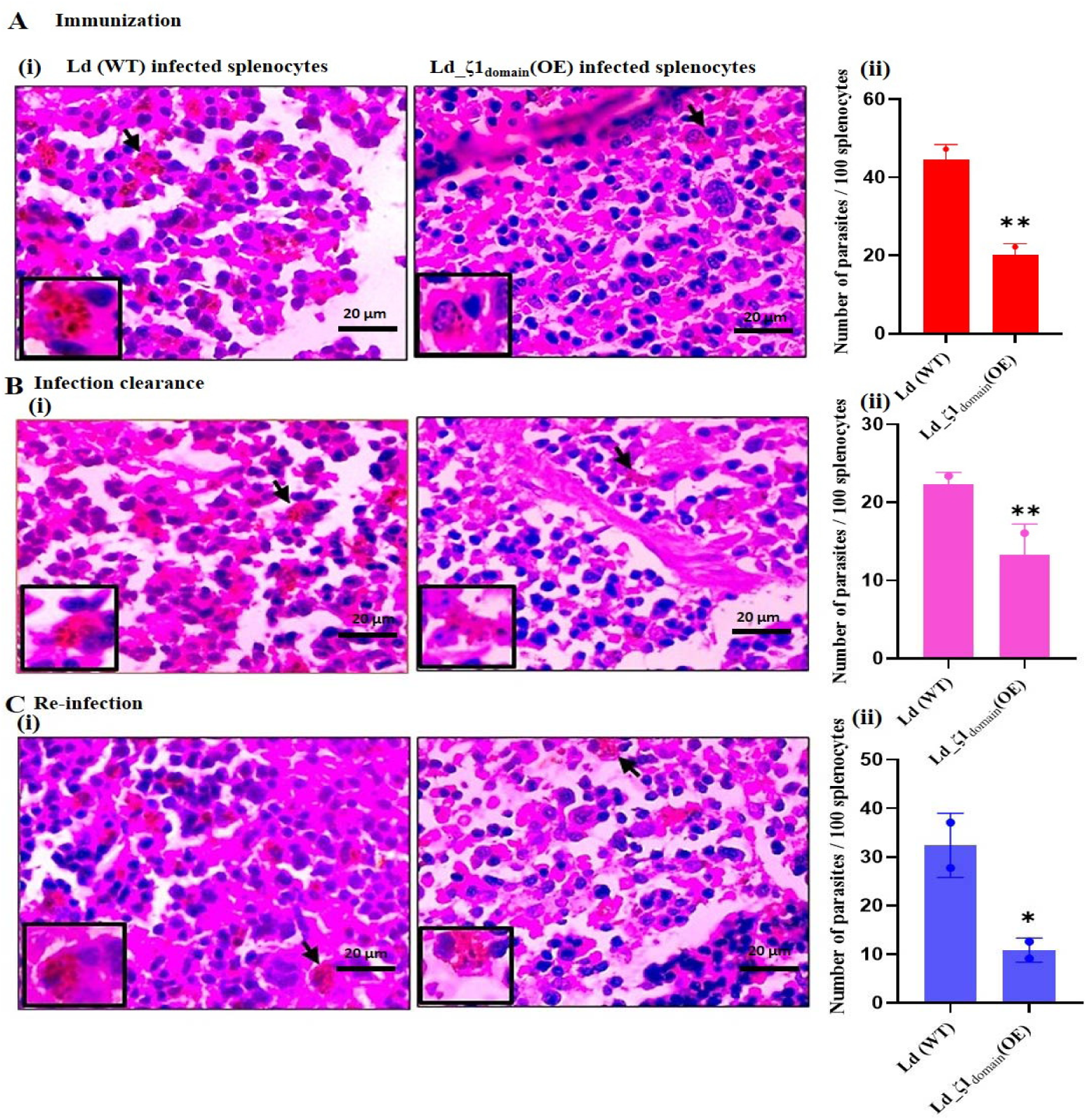
Photomicrographs showing splenic parasite burden in all groups following immunization **A(i)**, infection clearance **B(i)** and infectious challenge **C(i)** with *L. donovani*. Bar: 20 μm. Splenic tissues harboring a larger number of amastigotes (shown by arrowhead). Parasitemia was quantified by determining the number of parasites per 100 splenocytes **A(ii), B(ii) & C(ii)**.

Furthermore, hepatomegaly is the another key indicator of visceralization in leishmaniasis, particularly for diagnosing VL [21]. The preponderance of hepatic granulomas displayed an immature phenotype, characterized by a substantial presence of amastigotes in wild type infected mice **[Figure 6A (i)]**. Parasitemia of hepatic tissues exhibited a notable reduction by 50-60% in Ld_ζ1_domain_ (OE) infected mice in comparison to wild-type counterparts **[Figure 6A (ii)]**. After amphotericin treatment, parasite proliferation in both LdWT and Ld_ζ1_domain_ (OE) infected hepatocytes decreased **[Figure 6B (i) & (ii)]**. After reinfection, Ld_ζ1_domain_ (OE) immunized mice showed heightened immune response, resulting in a considerably decreased parasite load in the liver compared to LdWT control mice **[Figure 6C (i) &(ii)]**. Parasitemia quantification entailed plotting the count of amastigotes per 100 hepatocytes.

**Figure 6:**
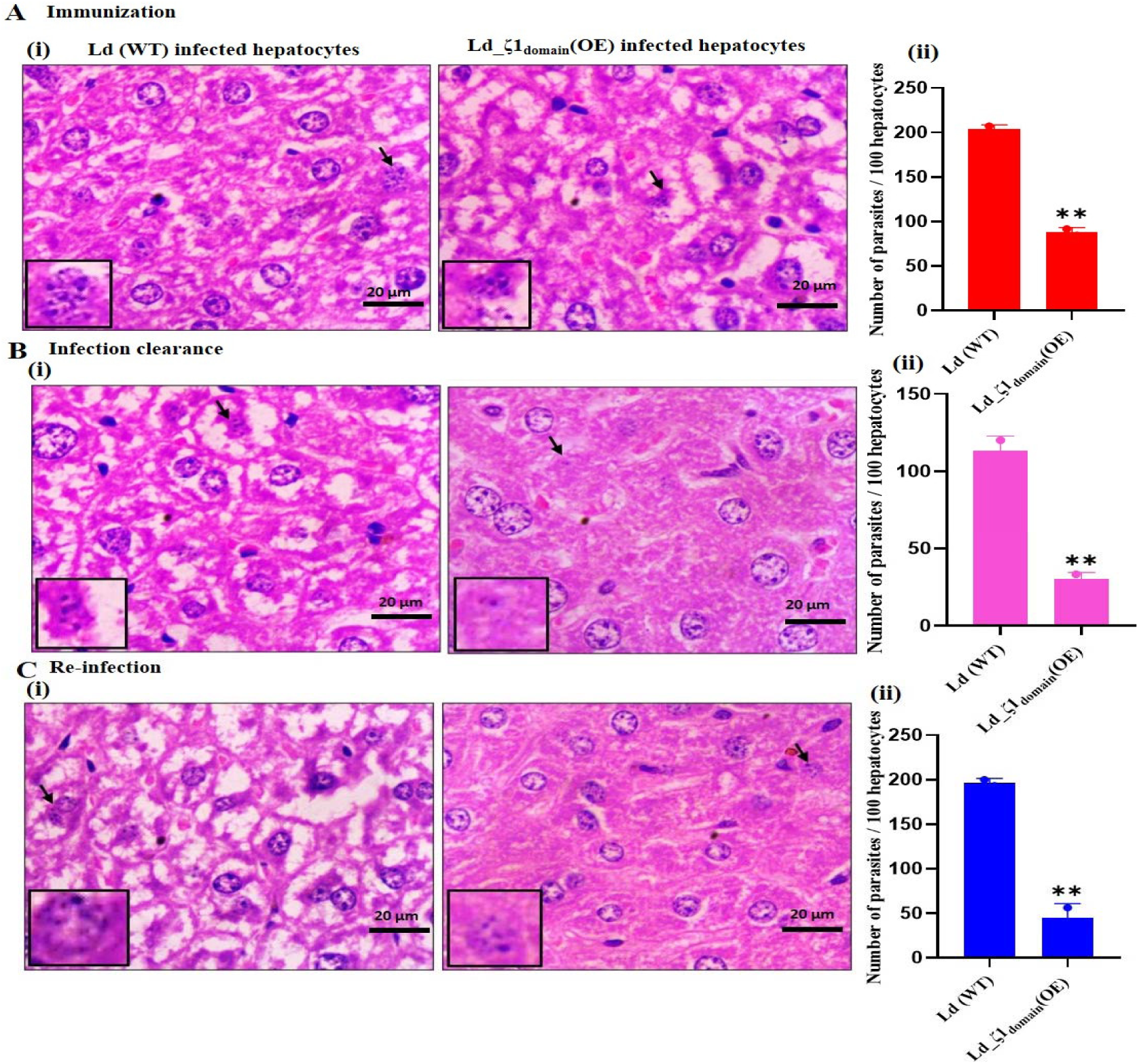
& Photomicrographs showing hepatic parasite burden in all groups following immunization **A(i)**, infection clearance **B(i)** and infectious challenge **C(i)** with *L. donovani*. Bar: 20 μm. Parasitemia was quantified by determining the number of parasites per 100 hepatocytes. Kupffer cell harboring a larger number of amastigotes (shown by arrowhead) **A(ii), B(ii) & C(ii)**.

### 3.6 Enhanced proinflammatory response and immune protection in Ld_**ζ**1_domain_ over-expressor immunized mice against *Leishmania* infection

Analysis of sera from immunized challenged mice revealed significantly increased *Leishmania*-specific IgG production in Ld_ζ1_domain_ over-expressor challenged mice compared to naive groups as depicted by ELISA **[Figure 7A]**. LdWT-infected mice exhibited heightened IgG antibody titres but significantly less than Ld_ζ1_domain_ (OE) challenged mice. This escalation serves to bolster the protective efficacy elicited by immunization with Ld_ζ1_domain_ (OE).

**Figure 7:**
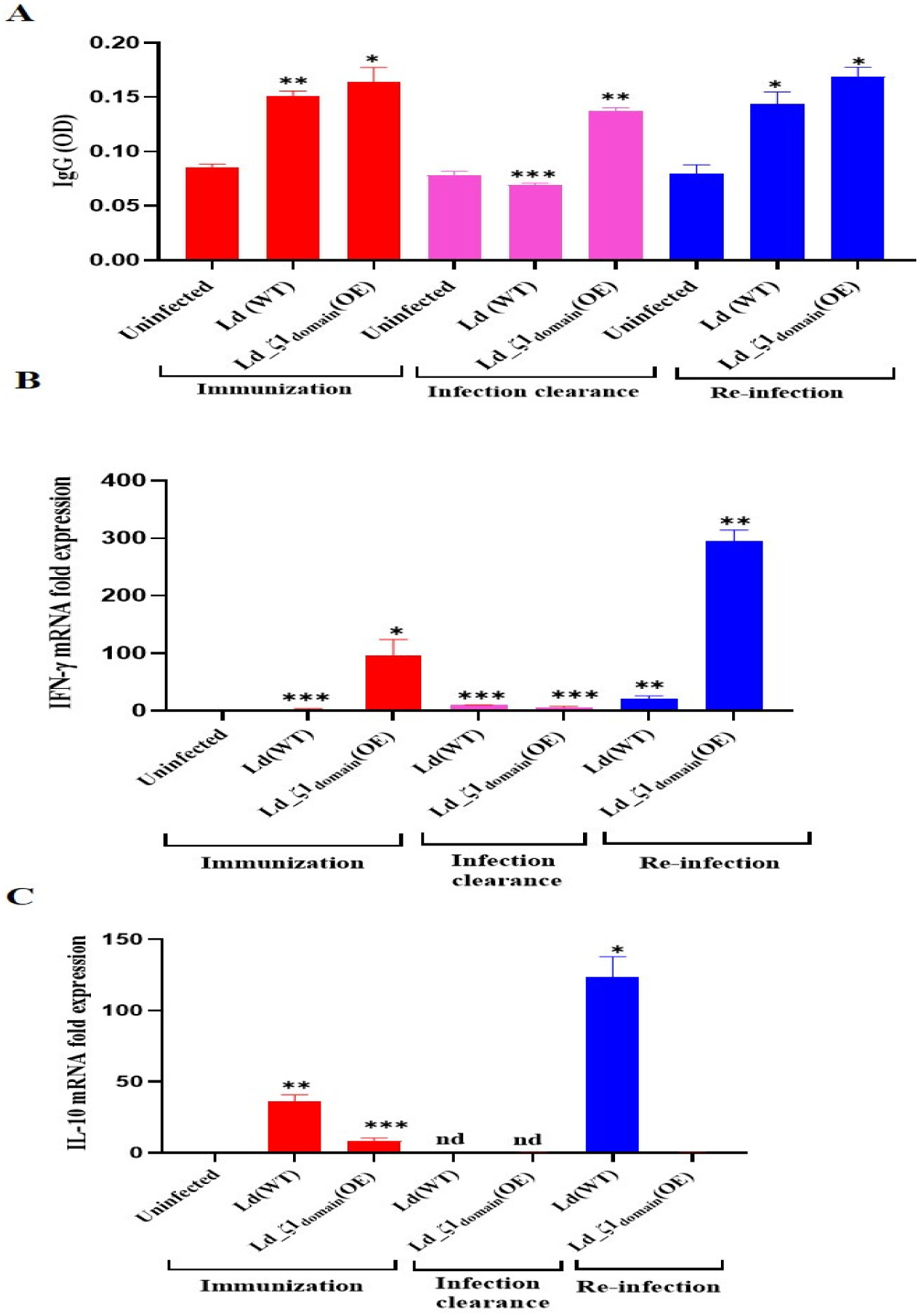
**Humoral and cellular immune responses induced in mice by immunization with Ld_**ζ**1_domain_**. **A**) Anti-*Leishmania* IgG serum levels in mice immunised with Ld_ζ1_domain_ (OE) and infected with *L. donovani*. **B** & **C**) Cytokine mRNA expression (IFN-γ and IL-10) levels in splenic tissues of mice immunised with Ld_ζ1_domain_ (OE) and infected with *L. donovani*. The data are presented as mean ± S.D. of four mice per group. Unpaired two-tailed Student’s t-test was performed to compare statistical significance at each time point. A P-value < 0.05 was considered significant. In all panels * represents P ≤ 0.05, ** represents P ≤ 0.01, and *** represents P ≤ 0.001.

Quantitative RT-qPCR assessed cytokine expression in spleen, focusing on IFN-γ **[Figure 7B]** and IL-10 **[Figure 7C]**. Vaccinated mice exhibited elevated IFN-γ and reduced IL-10 levels, indicating a pro-inflammatory shift. Elevated pro-inflammatory cytokines are associated with leishmaniasis resistance, while increased anti-inflammatory cytokines denote disease progression. IFN-γ to IL-10 ratio served as an immunization success indicator [22]. The observed alterations in spleen correlate with decreased mRNA expression of key pro-inflammatory and anti-inflammatory cytokines implicated in the immune response to *Leishmania*, including IFN-γ, TNF-α, and IL-10.

Furthermore, macrophage-derived nitric oxide (NO) is synthesized by iNOS (Inducible nitric oxide synthase) and acts as an antimicrobial agent. iNOS is triggered by pro-inflammatory cytokines such as interferon-gamma (IFN-γ). In VL-infected macrophages also, iNOS defends against *L. donovani* through its leishmanicidal NO molecule [23]. We observed elevated nitric oxide (NO) levels in Ld_ζ1_domain_ (OE) immunized mice at all time points. Notably, NO levels were significantly higher in reinfected mice at the 10^th^ week post-immunization, indicating a stronger immune response **[Figure 8A]**.

**Figure 8.**
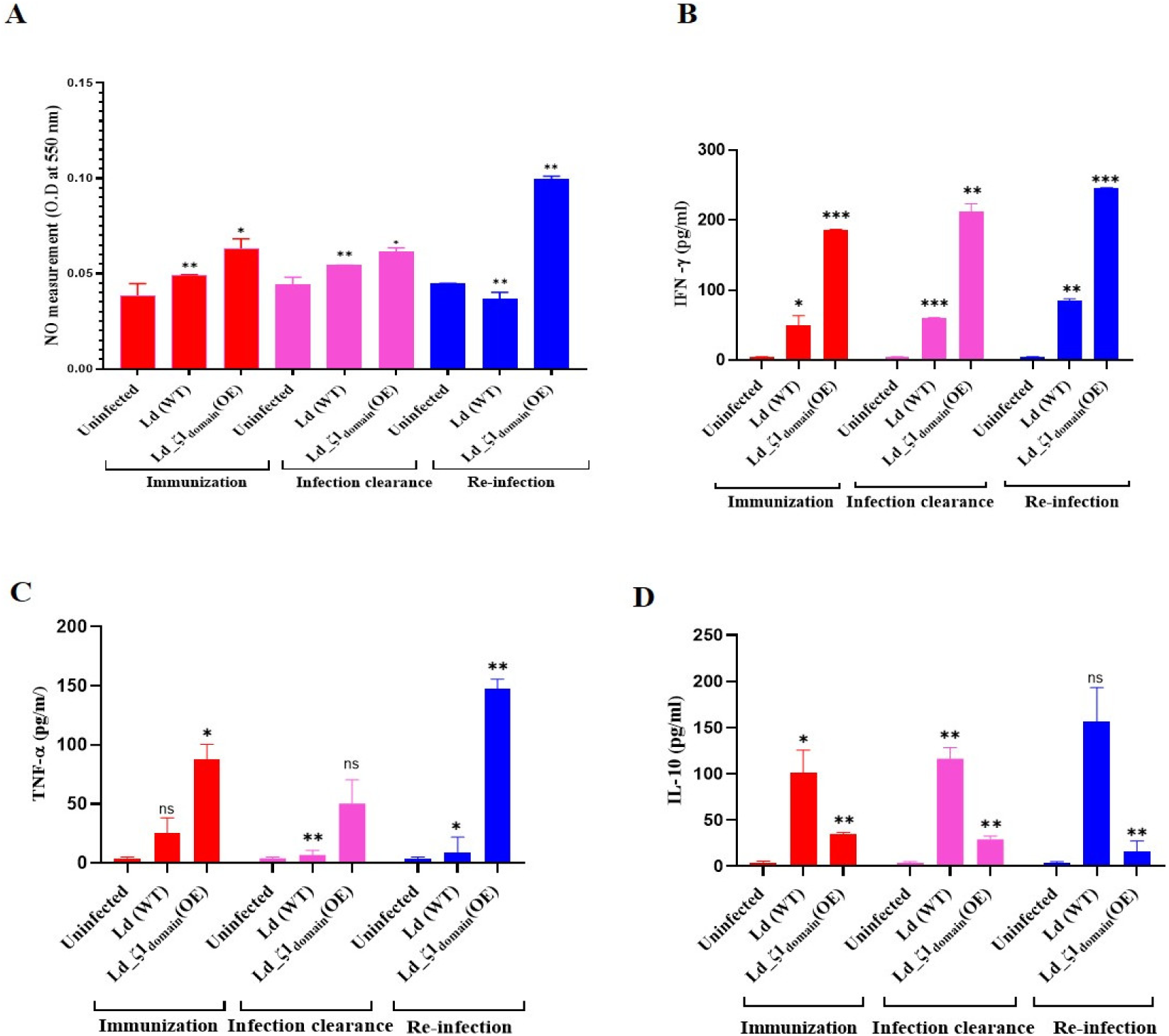
*Leishmania* Ag–stimulated cytokine profiles in splenocytes culture supernatants from Ld_ζ**1_domain_ (OE) immunized and LdWT challenged mice**. The immunized mice were euthanized, splenocytes were isolated, plated aseptically. (2×10^5^ cells/well), and stimulated with *Leishmania* FTAg for 48 h. **A**) NO levels in splenocytes isolated from immunized and non-immunized mice and challenged *in vitro* with *Leishmania* antigen by using the Griess Reaction Kit. Concentrations of pro-inflammatory cytokines IFN- γ (**B**), TNF-α (**C**) and IL-10 (**D**). The data are presented as mean ± S.D. of four mice per group. Unpaired two-tailed student’s t-test was performed to compare statistical significance at each time point. A P-value < 0.05 was considered significant. In all panels * represents P ≤ 0.05, ** represents P ≤ 0.01, and *** represents P ≤ 0.001.

Sandwich ELISA also confirmed higher levels of pro-inflammatory IFN-γ, TNF-α, and low level of anti-inflammatory cytokine IL-10 in stimulated splenocyte culture supernatants, leading to parasite clearance. **[Figure 8B, C & D]**.

## 4. Discussion

Toxin-antitoxin (TA) systems are common in bacterial genomes and promote survival against host defences [24].Our study explored a innovatory system in which *Leishmania* organisms are genetically modified to express a domain of “suicide-cum-attenuation” zeta toxin. This engineered gene was designed to weaken the parasites and simultaneously to reduce the virulence, providing an approach to controlling *Leishmania* infections more effectively.

Previously, a novel efficient regulatory expression system has been introduced in *Trypanosoma cruzi* which was based on the insertion of an inducible detrimental and toxin gene like degradation domain based on the *Escherichia coli* dihydrofolate reductase (ecDHFR) into the *T. cruzi* [25]. This DHFR degradation domain (DDD) was maintained by trimethoprim-lactate and degraded in the absence of induction agents. Amastigotes expressing one of the stimulated proteins of GFP-DDDHA, Alpha-toxin-DDDHA, Cecropin ADDDHA, or DDDHA were instigated to undergo a self-annihilation process within the host cells. In *in vivo* mouse models, these strains were found to be weakened and significantly protected mice against deadly infection with wild type strains. Several research have shown that various toxins were expressed in *L. mexicana* employing a recently created inducible protein stabilization system [26,27].Furthermore a novel suicidal xenotransgenic *Leishmania amazonensis* line was developed by episomally expressing δ-aminolevulinate dehydratase (ALAD) and porphobilinogen deaminase (PBGD) enzymes from the heme-biosynthetic pathway. This modification led to selective porphyrin accumulation in the parasite, enabling its photolysis to selectively destroy it within macrophage phagolysosomes. The transgenic *L. amazonensis* mutants showed potential as a safe, suicidal vaccine candidate by enhancing antigen presentation and T-cell immune responses [28].

In accordance with the abovementioned studies, we proposed designing a live attenuated vaccination approach by modifying the parasites by overexpressing its own inherited zeta toxin. This technique leverages the toxin’s elevated immunogenicity to promote a significant immune response, while attenuating the pathogen’s virulence ensures vaccination effectivity. In this direction, we cloned and overexpressed the Ld_ζ1_domain_ in *Leishmania* promastigotes using a GFP-tagged vector and confirmed the enhanced expression of zeta toxin in over-expressor by western blot and RT-PCR [**Figure 1C & 1E]**. We examined the growth dynamics of both promastigotes and amastigotes. Ld_ζ1_domain_-overexpressing and WT parasites grew similarly during the early log phase. However, starting with day 5 of the exponential development phase, Ld_ζ1_domain_ (OE) amastigotes displayed a considerable growth delay compared to the WT. Overexpression of the Ld_ζ1_domain_ limits amastigote formation while promastigote growth was unaffected **[Figure 2A]**.

We assessed Ld_ζ1_domain_ (OE) promastigote infectivity via giemsa staining, finding a 30-40% reduction in infected macrophages and a 75-85% decrease in intracellular parasites at 24 and 48 h compared to WT infections **[Figure 2B, 2C & 2D]**. Quantitative RT-PCR showed a significant reduction in infection in Ld_ζ1_domain_ (OE) infected THP-1 cells **[Figure 3A]**. This indicates that Ld_ζ1_domain_ overexpression significantly reduces parasite survival, replication, and infection rates. *In vitro* studies showed that in THP-1 cells infected with Ld_ζ1_domain_ (OE) versus LdWT parasites, TNF-α and IFN-γ levels were significantly higher, and IL-10 levels were lower after 24 h **[Figure 3B, 3C & 3D]**. This led to increased IFN-γ/IL-10 and TNF-α/IL-10 ratios, indicating improved parasite clearance and protection **[Figure 3E & 3F]**.

Next, we immunized mice with Ld_ζ1_domain_ (OE) and evaluated their protection against virulent *L. donovani*. Splenomegaly and hepatology as key VL symptoms were monitored. Non-immunized mice exhibited significant splenic enlargement while Ld_ζ1_domain_ (OE) immunized mice had spleen sizes similar to uninfected controls **[Figure 4B]**. RT-PCR analysis showed a sixfold higher parasite burden in LdWT infected mice compared to Ld_ζ1_domain_ (OE) immunized mice shortly after immunization. This higher parasitemia persisted in LdWT infected mice through the 7^th^ and 10^th^ weeks post-immunization **[Figure 4C]**.

Histopathology showed preserved splenic architecture with minor disarray in infected mice. LdWT infected splenocytes had higher infection rates post-immunization, while Ld_ζ1_domain_ over-expressor infected splenocytes had a 50% reduction **[Figure 5A]**. Amphotericin treatment reduced infections in both groups **[Figure 5B]**. Hepatomegaly showed immature hepatic granulomas with high amastigote levels in LdWT infected mice. Parasitemia in hepatic tissues was 50-60% lower in Ld_ζ1_domain_ (OE) infected mice, linked to immune response maturation **[Figure 6A]**.

Furthermore, humoral response pattern was also upregulated as evident by elevated anti-*Leishmania* IgG levels which clearly indicates a Th1 response [**Figure 7A]**. Immunized mice with Ld_ζ1_domain_ (OE) showed significantly higher *Leishmania* specific IgG [**Figure 7A]** and NO levels **[Figure 8A]** than naive and LdWT infected mice, suggesting a strong pro-inflammatory response. RT-qPCR analysis of spleen cytokines revealed increased IFN-γ and decreased IL-10 in vaccinated mice, indicating a pro-inflammatory shift [**Figure 7B & 7C]**. High pro-inflammatory cytokines (IFN-γ, TNF-α) in mice correlate with resistance to disease, while high anti-inflammatory cytokines (IL-10) indicates disease progression. The IFN-γ to IL-10 ratio marked immunization success. These findings were supported by sandwich ELISA measurements of IFN-γ, TNF-α, and IL-10 in splenocyte culture supernatants **[Figure 8B, 8C & 8D]**.

Collectively, the results indicated that the overexpression of the zeta toxin caused regulated attenuation of the parasites, reducing their pathogenicity while retained their immunogenic characteristics. Our study demonstrated the protective effectiveness, immunogenicity, and proliferation of the zeta over-expressor in response to *Leishmania* challenge *in vitro* and *in vivo* model **[Graphical representation]**. This exploratory pilot study stipulated that Ld_ζ1_domain_ (OE) parasites are potential candidates for the generation of an attenuated vaccine against leishmaniasis.

## 5. Conclusion

In conclusion, our findings showed that overexpression of the zeta toxin in *Leishmania* parasites reduced pathogenicity while retaining immunogenicity. *In vitro*, this genetic alteration lowered parasite survival, replication, and infection rates, but *in vivo* it enhanced cellular and humoral immune responses and provided protection against virulent *L. donovani*. The Ld_ζ1_domain_ OE parasites increased pro-inflammatory cytokines (IFN-γ and TNF-α) while decreasing anti-inflammatory cytokines (IL-10), resulting in a Th1 biased immune response and better parasite clearance. This data indicated that the Ld_ζ1_domain_ (OE) parasites are prospective candidates for developing a live attenuated vaccination against leishmaniasis.

## Author Contributions

**Shailja Singh**: Writing – original draft, Conceptualization, Methodology, Supervision, Resources, Data curation, Validation & Funding acquisition. **Ruby Bansal**: Writing – review & editing, Writing – original draft, Conceptualization, Data curation & Methodology. **Sadat Shafi:** Methodology, Animal studies & Formal analysis. **Prachi Garg**: Visualization, Protein purification & Western blots. **Aakriti Srivastava**: Methodology & Visualization. **Swati Garg**: Methodology, Software & Investigation**. Neha Jha:** Methodology & Investigation. **Jhalak Singhal**: Methodology & Project administration. **Gajala Deethamvali Ghouse Peer**: Methodology. **Ramendra Pati Pandey**: Formal analysis, Software & Investigation. **Subhajit Basu:** Validation, Software & Investigation.

